# Phenotypic variability of hydraulic residual conductance and its temperature sensitivity in *Abies alba* Mill

**DOI:** 10.64898/2026.01.22.700907

**Authors:** Stéphane Herbette, Stéphane Andanson, Alexandre Gonzalez, Chris J. Blackman, Julien Cartailler, Ludovic Martin, Hervé Cochard

## Abstract

Residual water losses after stomatal closure have recently been identified as key determinants of drought-induced hydraulic failure, particularly under heatwave conditions. However, little is known about the intraspecific variability of residual conductance (*g*_res_) and its thermal sensitivity. Here, we investigated the genetic and environmental sources of variation in *g*_res_ and its associated thermal parameters (phase transition temperature *T*_, and temperature sensitivities *Q*_10a_ and *Q*_10b_) in *Abies alba* Mill., together with vulnerability to xylem embolism (*P*_50_). Measurements were performed using the Drought-Box on seven French provenances grown in a common garden to assess genetic variability, and on trees growing across contrasting forest sites to quantify phenotypic plasticity. Seasonal dynamics and within-canopy microclimatic effects were also examined, and linked to needle biochemical traits.

Residual conductance exhibited a marked seasonal decline, with high values in newly formed needles followed by a stabilization from late summer to the following spring, closely tracking the accumulation of cuticular waxes. In contrast, Klason lignin content showed little seasonal variation. Difference between provenances was weak for all investigated parameters, suggesting strong constraints on these safety-related traits. By contrast, *g*_res_ showed significant environmental plasticity, with lower values at more climatically constrained sites, while thermal parameters and *P*_50_ remained relatively conserved.

Our results identify *g*_res_ as a developmentally dynamic and environmentally plastic trait in silver fir, potentially representing a key lever of acclimation to drought. Incorporating such variability into mechanistic models should improve predictions of tree vulnerability under future climates combining intensified droughts and heatwaves.

**Key message.:** Residual conductance in *Abies alba* is developmentally dynamic and environmentally plastic but genetically constrained, highlighting its key role in acclimation to drought and heatwave-driven hydraulic failure.

## Introduction

Massive tree mortality events have been observed worldwide due to an increase in the frequency of heat waves coupled to drought events (Allen et al. 2010; Choat et al. 2012; Hammond et al. 2022). Under drought conditions, increasing tension within the xylem can exceed a threshold value causing the formation of air-embolisms that block sap flow; and high temperatures combined with drought exacerbate the phenomenon (Cochard 2021). This so-called catastrophic hydraulic failure is a major cause of tree deaths during drought (Barigah et al. 2013; Anderegg et al. 2016; Mantova et al. 2022; Hammond et al. 2022). Plants reduce the risk of hydraulic failure during drought through different physiological mechanisms such as decreasing xylem vulnerability to embolism and maintaining favorable plant water status through stomatal closure and low residual transpiration (Martin-StPaul et al. 2017). Characterizing how different plant species modulate these traits will help in determining their risk of drought-induced mortality and is crucial for predicting how climate change will affect species and vegetation distribution worldwide (Anderegg 2015).

The determination of forest or tree mortality risk under drought conditions is also challenged by the ability of trees to adapt or to acclimate to changing conditions. Indeed, a wide range of structural and physiological adjustments based on phenotypic plasticity and genetic variability allow trees to adapt to environmental changes, such as those imposed through climate change (Sultan 2000; Nicotra et al. 2010). Several reports point to the importance of acclimation process in delaying the onset of mortality or hydraulic failure (Brodribb et al. 2020; Lemaire et al. 2021; Barigah et al. 2023; Andriantelomanana et al. 2024). Genetic variability could also be a key feature in forest adaptation to climate change. Replanting forest stands with genotypes of the same species that are better adapted to drought conditions is appealing, but little information is available on the genetic diversity of drought tolerance in tree species. A study performed on *Vitis* highlighted the potential of genetic variability in a set of hydraulic safety traits to protect elite genotypes from hydraulic failure under future climate scenarios (Dayer et al. 2022). These traits were quantified within and across *Vitis* species, then traits combinations were created in silico to identify tolerant trait syndromes. Such results on acclimation and genetic variability call for a more complete characterization of the within species variability of the most important hydraulic traits in tree species. For more than a decade, numerous studies have been carried out on the within-species variability of vulnerability to xylem embolism (Awad et al. 2010; Herbette et al. 2010; Lamy et al. 2011; Wortemann et al. 2011) and stomatal closure (Bartlett et al. 2014), as both traits were identified as key traits in drought tolerance (Choat et al. 2012; Delzon and Cochard 2014). However, when stomata are closed, plants continue to lose water through leaf cuticles, leaky stomata and bark lined with lenticels. Such residual water losses have been found to influence the kinetics of hydraulic failure (Blackman et al. 2016, 2019; Martin-StPaul et al. 2017). According to simulations using the mechanistic model ‘SurEau’ (Martin-StPaul et al. 2017; Cochard 2021), the residual leaf (*g*_min_) or shoot (*g*_res_) conductance has been identified as a key trait determining the onset of hydraulic failure. A recent experiment confirmed the importance of *g*_res_ in controlling the time to hydraulic failure for evergreen species (Blackman et al. 2023). Moreover, the role of residual conductance would be exacerbated under conditions of drought coupled with a heatwave (Cochard 2021). Indeed, *g*_res_ are sensitive to increasing temperature (Bueno et al. 2019; Billon et al. 2020). The temperature threshold where *g*_res_ markedly increases has been called the phase-transition temperature (*T*_p_), and corresponds to the temperature above which cuticular permeability increases as a result of changes in the structure of cuticular waxes (Riederer and Muller 2008). Cochard (2021) proposed to evaluate the temperature dependance of *g*_res_ by calculating *Q*_10a_ et *Q*_10b_, i.e. the slope of *g*_res_ as a function of temperature (*Q*_10_) below and above the *T*_p_, respectively. These *g*_res_-related traits need to be considered to improve prediction of the risk of hydraulic failure during heatwaves periods (Cochard 2021). The importance of *g*_res_ and its *T*_p_, requires further study of their within-species variability in order to improve our knowledge of trees’ ability to acclimate or adapt to severe drought conditions. These traits vary across species and *g*_res_ seems to be correlated with environmental conditions such as rainfall in some groups of plants (Duursma et al. 2019; Blackman et al. 2023). Investigations on within-species variability of *g*_res_ are scarce (Hasanuzzaman et al. 2017; Lemaire et al. 2021), and to our knowledge, they have not been undertaken for *T*_p_ , *Q*_10a_ or *Q*_10b_. This could be explained by the lack of a relevant method to screen difference between genotypes for these traits (Hasanuzzaman et al. 2023).

Silver fir (*Abies alba* Mill.) is an interesting species for investigating risk of drought-induced mortality in the context of climate change. It is a dominant resinous species in the humid mountainous regions of central and southern Europe, ranging from the Pyrenees to the Carpathians and from Italy to Poland. It is an important species contributing to biodiversity conservation and timber production (Walder et al. 2021), and it has some advantage such stand resistance to wind and insect attack in mixture with more sensitive species such as *Picea abies* (Wolf 2003). However, the ability of silver fir to cope with climate change is unclear. This species is considered well-adapted to moist conditions and poorly adapted to warm and summer drought conditions. Moreover, widescale drought-induced tree decline and dieback have been observed for several years for silver fir (Camarero et al. 2011; Cailleret et al. 2014). On the contrary, paleoecological studies reported its former presence in warmer and dryer conditions (Tinner et al. 2013), while the current absence under such conditions would be due to the long-lasting human pressure on this fire- and browsing-sensitive species (Carcaillet and Muller 2005). Moreover, Silver fir is expected to be more drought-resistant compared to its main co-occuring species, including Fagus sylvatica or Picea abies, thanks to its deeper root system (Magh et al. 2017), and the better resistance of its growth to drought episodes (Walder et al. 2021). To get additional clues on the ability of silver fir to cope with increasing drought events, we need to investigate its ability to adapt or acclimate through genetic variability or phenotypic plasticity for key drought-resistant traits.

In this study, we conducted an investigation of the variability of *g*_res_ and its related parameters (*T*_p_ , *Q*_10a_ or *Q*_10b_ ) in Silver fir, as well as the variability of vulnerability to xylem embolism, another key trait of hydraulic safety. To evaluate the genetic variability among French provenances, we measured these traits on seven provenances grown in a common garden. To investigate the phenotypic plasticity, we measured the same traits on trees grown in different forest sites contrasting in altitude and rainfall, and according to sunlight exposure, as this latter has been shown to impact hydraulic safety traits such as vulnerability to embolism and stomatal closure (Herbette et al. 2010; Gerardin et al. 2018). We also studied the *g*_res_ across the season and we investigated the underlying biochemical and structural determinants of this marked seasonal evolution. For accurate measurements of *g*_res_ and its related parameters, we used a new automated tool, the DroughtBox (Billon et al. 2020). It consists of a programmable climatically controlled chamber in which branches dehydrate and changes in the mass recorded. It is a reliable tool for investigating within species variability for g_res_ , *T*_p_ , *Q*_10a_ or *Q*_10b_.

## Materials and methods

### Plant material and sites

The studies were conducted using shoots sampled from silver fir trees grown in different and contrasting environment according to the questions addressed. The forest sites and fir provenances are described in Table 1.

**Table 1.**
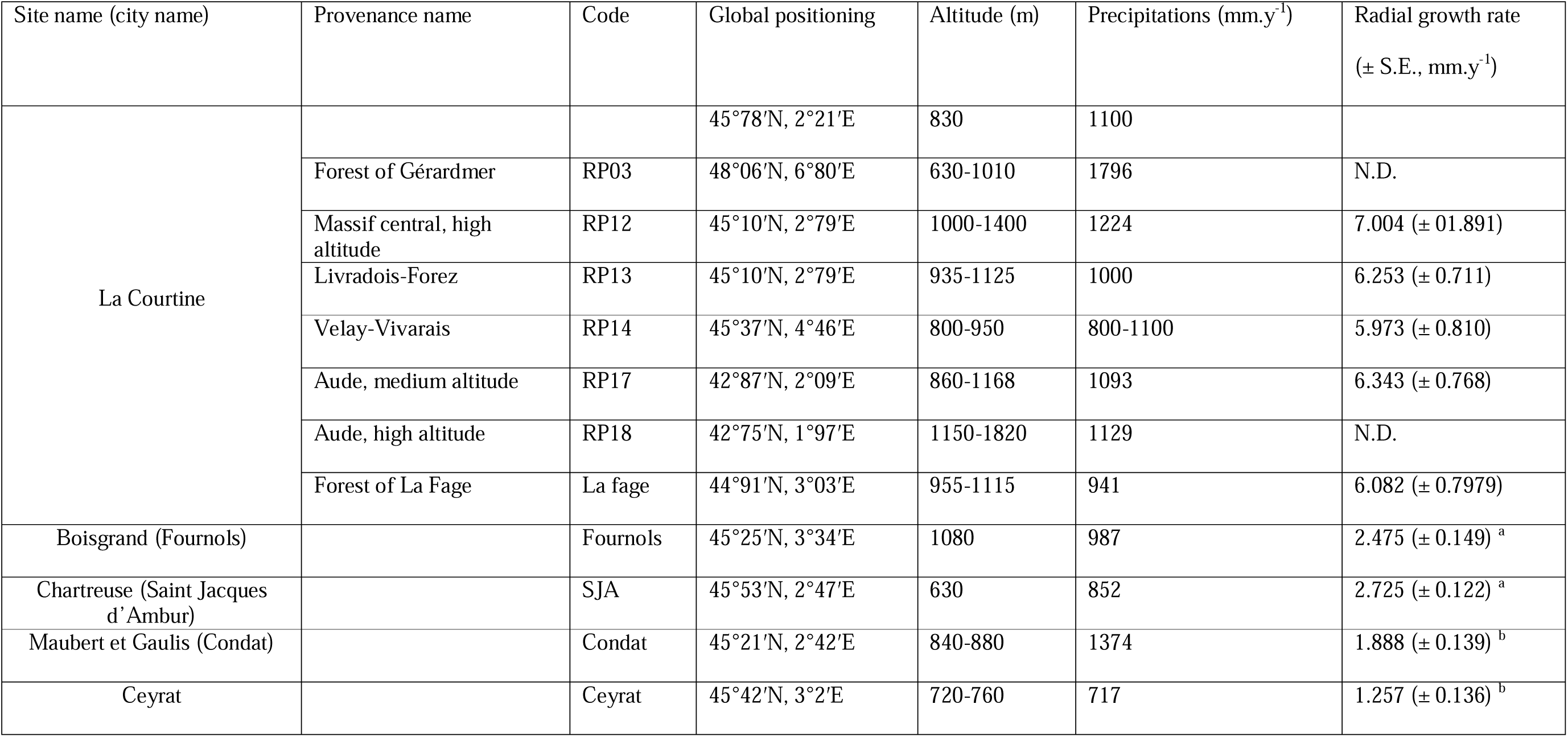
Characteristics of study sites and of the seven Fir provenances evaluated in this study. The global positioning was given for the origin location of each Fir provenance as well as for the common garden where they were planted (La Courtine). The radial growth rate was measured on 8 to 16 trees and analyzed for the years 2015 to 2020 for Fournols, SJA, Condat and Ceyrat sites. Data are mean values (±S.E) and different letters indicate significant differences between the four forest sites. The radial growth was also measured on 30 to 38 trees and analyzed for the years 2011 to 2018 for 5 Fir provenances grown in the forest site of La Courtine. Data are mean values and no significant difference was found between these provenances.

| Site name (city name) | Provenance name | Code | Global positioning | Altitude (m) | Precipitations (mm.y<sup>-1</sup>) | Radial growth rate<br>(± S.E., mm.y<sup>-1</sup>) |
| --- | --- | --- | --- | --- | --- | --- |
| La Courtine |  |  | 45°78'N, 2°21'E | 830 | 1100 |  |
|  | Forest of Gérardmer | RP03 | 48°06'N, 6°80'E | 630-1010 | 1796 | N.D. |
| | Massif central, high altitude | RP12 | 45°10'N, 2°79'E | 1000-1400 | 1224 | 7.004 (± 01.891) |
| | Livradois-Forez | RP13 | 45°10'N, 2°79'E | 935-1125 | 1000 | 6.253 (± 0.711) |
| | Velay-Vivarais | RP14 | 45°37'N, 4°46'E | 800-950 | 800-1100 | 5.973 (± 0.810) |
| | Aude, medium altitude | RP17 | 42°87'N, 2°09'E | 860-1168 | 1093 | 6.343 (± 0.768) |
|  | Aude, high altitude | RP18 | 42°75'N, 1°97'E | 1150-1820 | 1129 | N.D. |
| | Forest of La Fage | La fage | 44°91'N, 3°03'E | 955-1115 | 941 | 6.082 (± 0.7979) |
| Boisgrand (Fournols) | | Fournols | 45°25'N, 3°34'E | 1080 | 987 | 2.475 (± 0.149) <sup>a</sup> |
| Chartreuse (Saint Jacques d'Ambur) | | SJA | 45°53'N, 2°47'E | 630 | 852 | 2.725 (± 0.122) <sup>a</sup> |
| Maubert et Gaulis (Condat) | | Condat | 45°21'N, 2°42'E | 840-880 | 1374 | 1.888 (± 0.139) <sup>b</sup> |
| Ceyrat | | Ceyrat | 45°42'N, 3°2'E | 720-760 | 717 | 1.257 (± 0.136) <sup>b</sup> |

The common garden experiment was located at La Courtine, in central France. The site experiences a montane climate highly favorable for the growth and development of silver fir (*Abies alba* Mill.), and the plantation was established in 1995 and hosts several French provenances of the species grown under homogeneous environmental conditions. Seven provenances originating from contrasting climatic and altitudinal regions across France were selected for this study, spanning a wide latitudinal, altitudinal (ca. 630–1820 m a.s.l.) and precipitation gradient (800–1800 mm.yr_¹ at the site of origin). This design allowed us to assess the extent of genetic variability in hydraulic safety traits independently of environmental effects. Growth measurements revealed no significant differences in radial growth among provenances at La Courtine (Table 1), indicating comparable tree vigor and supporting the use of this common garden to investigate genetic variability in hydraulic traits.

To assess the influence of environmental conditions on hydraulic traits, silver fir trees were also sampled in several forest stands. The selected forest sites are located in central France and span a gradient of climatic favorability for silver fir, mainly driven by differences in altitude and annual precipitation (Table 1). Radial growth measurements revealed significant differences among forest sites, supporting the use of these stands as a natural gradient of environmental constraints (Table 1). These contrasting conditions were used to quantify the extent of phenotypic plasticity in hydraulic traits, in comparison with trees grown under the favorable and homogeneous conditions of the common garden.

Seasonal and microclimatic effects on residual conductance and associated traits were investigated at a single low-altitude forest site located in the commune of Neuville, central France (45.7519° N, 3.4556° E; 465 m a.s.l.). Seasonal variability was assessed on three individual silver fir trees by repeatedly measuring residual conductance (*g*_res_) and related biochemical traits on the same cohort of needles produced in 2022. Measurements were conducted on 19 May, 8 July, 27 August and 16 October 2022, and on 24 January, 15 March, 4 May and 23 June 2023, thus covering a full annual growth and hardening cycle. In parallel, within-canopy microclimatic effects were examined by comparing sun-exposed upper branches and shade-exposed lower branches sampled on the same individuals. This combined design allowed disentangling seasonal effects from light-related microenvironmental variation while minimizing confounding influences of site or regional climate.

### Pressure–volume curves and turgor loss point determination

Pressure–volume (PV) curves were constructed to determine the leaf turgor loss point (Ψ_TLP_) of silver fir. Sun-exposed branches were collected and immediately transported to the laboratory in humid, dark conditions to minimize dehydration. Branches with needles attached were rehydrated overnight by immersing the cut ends in distilled water for full saturation. Leaf water potential (Ψ_leaf_) was measured using a Scholander-type pressure chamber (PMS, Corvallis, OR, USA), while fresh mass was recorded using a precision balance (Mettler AE 260, DeltaRange®; Mettler Toledo, Columbus, OH, USA). Following full hydration, branches were allowed to dehydrate at room temperature. During dehydration, Ψ_leaf_ and leaf mass were measured repeatedly. After completion of the PV measurements, needles were oven-dried at 70 °C for 3 days to determine dry mass. Pressure–volume curve parameters, including osmotic potential at full turgor, Ψ_TLP_, relative water content at turgor loss point (*RWC*_TLP_), and bulk modulus of elasticity (ε), were calculated according to standard procedures (Sack et al. 2010).

### Residual conductance, *T*_p_, *Q*_10a_ and *Q*_10b_

The minimum residual conductance (*g*_res_, mmol.m^-2^.s^-1^) and its phase transition temperature (*T*_p_) were determined using five to six small branches, per experiment, dried inside a temperature- and relative humidity-controlled Drought-box (Billon et al. 2020). The branches, 30 to 40 cm long, were hydrated overnight, air-cut and the cut end sealed with melted paraffin. The length (mm), basal diameter (mm) and saturated mass (g) of each branch were measured before being suspended from one of the eight strain gauges inside the Drought-box. For the *g*_res_ vs. season experiment, climatic conditions were set at 30°C and 40% relative humidity, and branches were allowed to dehydrate, with changes in mass representing water loss from the branch. Stomatal closure occurred after a period ranging from 1 to 5 hours. After 24 hours in the Drought-box, the branches were removed and the leaves and stems were oven-dried at 70°C to obtain the dry mass. The *g*_res_ was determined from water loss data beyond the point of stomatal closure. To do this, water loss rates were normalized by the vapor pressure deficit (VPD; KPa) recorded inside the Drought-box and the surface area of the main stem (represented by a cylinder) and the projected leaf area (*LA*). The *LA* was calculated by multiplying the total dry weight of the leaves on each branch by the average *SLA* for the species.

In the other experiments, not only was *g*_res_ measured, but also the phase transition temperature (*T*_p_, °C), which defines the temperature point at which *g*_res_ increases markedly over a range of temperatures, was determined on a separate set of branches taken from plants in each condition and dried in the drought-box (Billon et al. 2020). In these experiments, branches were dried under six progressive temperature increases (30, 35, 40, 45, 50 and 55°C), with simultaneous decreases in relative humidity (37, 24, 17, 13, 11 and 10% for the curtain wall samples and 40, 30, 23, 18, 14 and 11% for the comparison of the different forests). The *g*_res_ was calculated at each temperature step as described above. From these data, we calculated *T*_p_ for each provenance using an Arrhenius diagram as described in Bueno et al. (2019), while *Q*_10a_ and *Q*_10b_ were calculated as the *Q*_10_ of the relationship between *g*_res_ and temperature below and above the *T*_p_ value, respectively.

### Xylem vulnerability to embolism

Vulnerability to embolism was measured on branches located at the top of the trees and exposed to sunlight when no specified. Branches of 0.4–0.5 m long were harvested from 16 individuals for comparisons among forest sites and 11 individuals for comparisons among provenances or between sun-exposure treatments. Branches were wrapped in humid paper inside plastic bags, brought to the laboratory and stored at 4 °C until measurements in the following days, up to 1 week after sampling. We cut stem segments of 0.28 m-long just before the measurement of their vulnerability to cavitation using the Cavitron technique (Cochard et al. 2005). We measured the loss of hydraulic conductance of the stem segment while the centrifugal force generated an increasing negative xylem pressure. The plot of the percent loss of xylem conductance (*PLC*) versus the xylem pressure represents the vulnerability curve. First, we set the xylem pressure to a reference pressure (−0.5 or −1.0 MPa) and we determined the maximal conductance (*k*_max_) of the sample. Then, we set the xylem pressure to a more negative pressure for 2 min, and we determined the new conductance (*k*). The procedure was repeated for more negative pressures (with −0.50 MPa increments) until *PLC* reached at least 90%. We generated vulnerability curves for each branch by fitting the data with exponential–sigmoidal function (Pammenter and Van der Willigen 1998):

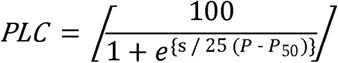

where *P*_50_ is the pressure causing 50% loss of conductance and *s* is the slope value at this point.

### Needles cuticular wax mass

Samples were frozen at -80°C immediately after sampling, then thawed and processed all at the same time. Wax was extracted by solubilization in chloroform (≥ 99.5%, Sigma-Aldrich) according to the protocol proposed by Bueno et al. (2019) with minor adaptations to the studied species. Briefly, 500 mg of needles per sample were immersed successively in two 20 mL chloroform bath for 5 min each. The resulting extracts were pooled and evaporated under a nitrogen stream. The mass of extracted wax was determined using an analytical balance (Mettler Toledo ME 204). Mass values ranged from 3.4 to 5.2 mg, with an associated compound standard uncertainty of 0.25 mg. The needles were then recovered and scanned for total surface area using Fidji software (ImageJ version 1.53t). Cuticular wax content was expressed on a surface basis (g.mm^-2^).

### Klason lignin rate

For each sample, 4 g of branches bearing fresh needles were placed at -80°C. Once all the samples had been collected, they were oven-dried at 50°C for 5 days. The needles were then separated from the branches and ground using an ultracentrifugal mill (Retsch ZM200, 0.75 mm ring sieve). The resulting powder was sieved with a sieve shaker (Retsch AS200, 75 µm + 500 µm sieve; 100 mm) for 5 min to collect particles ranging from 75 to 500 µm. Soluble compounds were removed by successive hot extractions. Briefly, 500 mg of dry matter were placed in heat-sealable bags (ANKOM F57, 25µm) and successively immersed in boiling 96% ethanol for 30 min, followed by boiling ultrapure water for 30 min. This extraction cycle was repeated seven times. The resulting cell wall residue was dried at 105°C for 16 h. Klason lignin content was determined according to the Tappi T222 om-88 standard. An aliquot of 300 mg of the dried residue was incubated with 5 mL of 72% sulfuric acid for 1 h at room temperature under continuous stirring with glass beads to hydrolyze cellulose and hemicelluloses. The sample was then diluted with 193.5 g ultrapure water to achieve a final sulfuric acid concentration of 3% and autoclaved at 120°C for 1 h. After cooling, the hydrolysate was filtered through a sintered glass filter (pore size of 48 mm). The residue was washed twice with 800 mL ultrapure water until neutral pH was reached. The filter containing the acid insoluble residue was dried at 105°C for 16 h and cooled in a desiccator. The dried residue, corresponding to Klason lignin, was weighted and expressed as a percentage of initial dry matter. Each sample was analyzed in duplicate.

### Statistical analysis

To investigate differences in physiological traits in relation to season, temperature, tree provenances, and environment, we tested for differences among groups. Normality of the data was assessed using the Shapiro–Wilk test (shapiro.test function in R, R Stats Package, v.4.4.2) and homoscedasticity of variances was evaluated using Levene’s test (leveneTest function, R CAR Package, v.3.1-3). If the data were normally distributed and homoscedasticity was verified, we used analysis of variance (ANOVA) (aov function, R Stats Package, v.4.4.2), followed by Tukey’s Honest Significant Difference (HSD) test (TukeyHSD function, R Stats Package, v.4.4.2) for post hoc pairwise comparisons. If normality or homoscedasticity was not verified, we performed a Kruskal-Wallis rank sum test (kruskal.test function, R Stats Package, v.4.4.2) as a non-parametric alternative. If a significant difference was detected (p < 0.05), we conducted Dunn’s Kruskal-Wallis multiple comparisons test (dunnTest function, R FSA Package, v.0.9.5) for post hoc analyses. All tests were performed with a significance level of α = 0.05.

## Results

### Seasonal variation of residual conductance (gres)

Residual conductance (*g*_res_) varied significantly over the annual cycle when measured on the same individuals and the same cohort of needles (Fig. 1). The highest values were recorded early in the growing season (June 2022), when current-year needles were fully expanded. Thereafter, *g*_res_ decreased markedly and reached lower values by mid-summer. From late summer through winter, *g*_res_ remained relatively stable, and similar values were observed again the following spring on one-year-old needles.

**Fig. 1.**
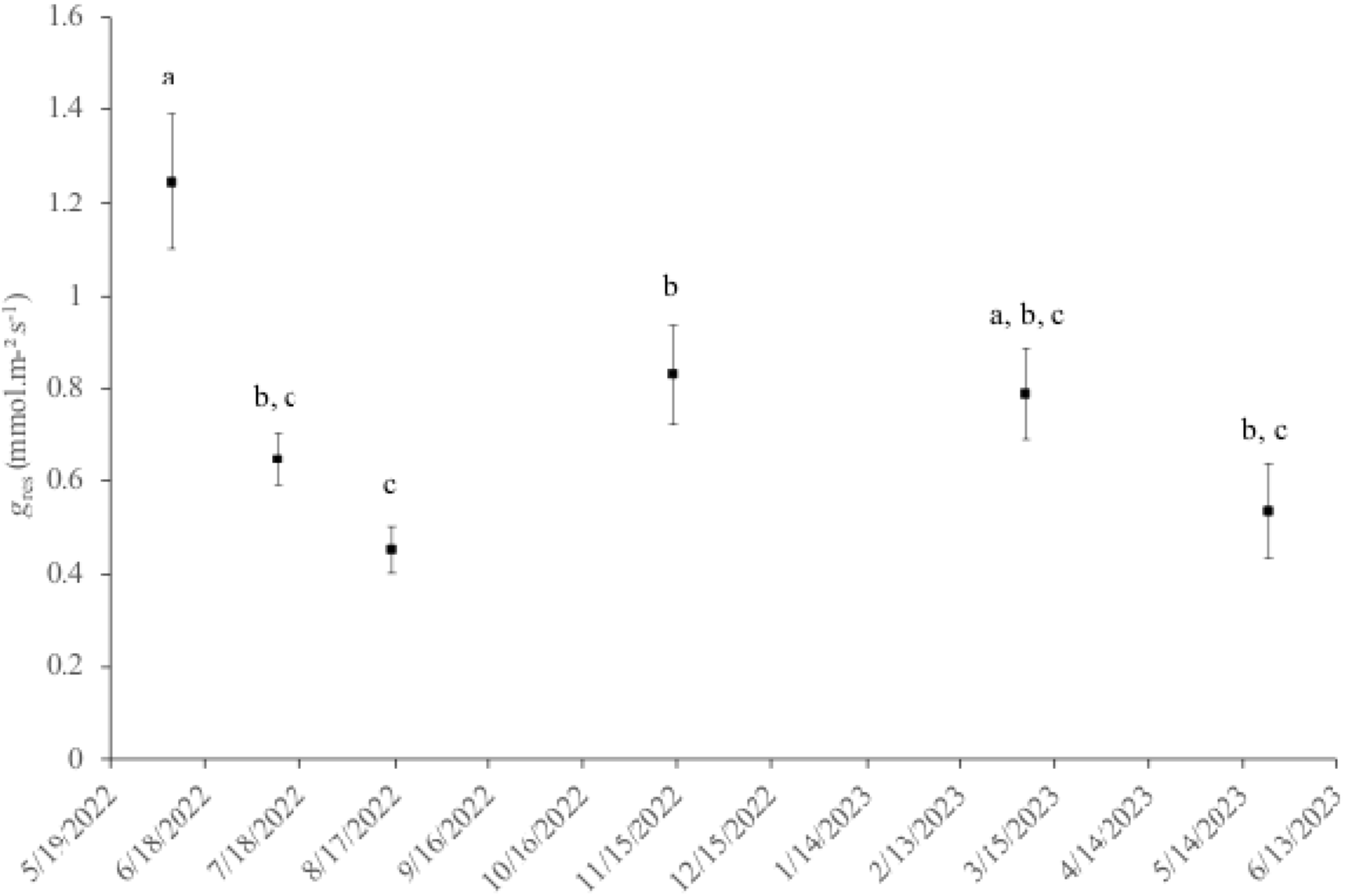
Evolution of *g*_res_ across the seasons. The individuals monitored are the same, as is the cohort of needles. The 06/07/2022 measurements are taken on the most recent, fully developed needles, while those taken on 05/22/2023 are 1 year old. Data are the mean values (± S.E.). Different letters indicate significant differences between sampling dates.

### Inter-provenance variability in *g*_res_ and hydraulic safety traits

Residual conductance measured across a range of temperatures did not show consistent differences among the seven provenances grown in the common garden (Fig. 2). Although some differences were detected at specific temperatures, no systematic provenance-specific pattern was observed. Accordingly, no significant differences were found among provenances for *T*_p_, *Q*_10a_, or *Q*_10b_ (Table 2). Vulnerability to xylem embolism (*P*_50_) also varied within a narrow range and did not differ significantly among provenances. When available, Ψ_TLP_ showed similarly limited variation.

**Fig. 2.**
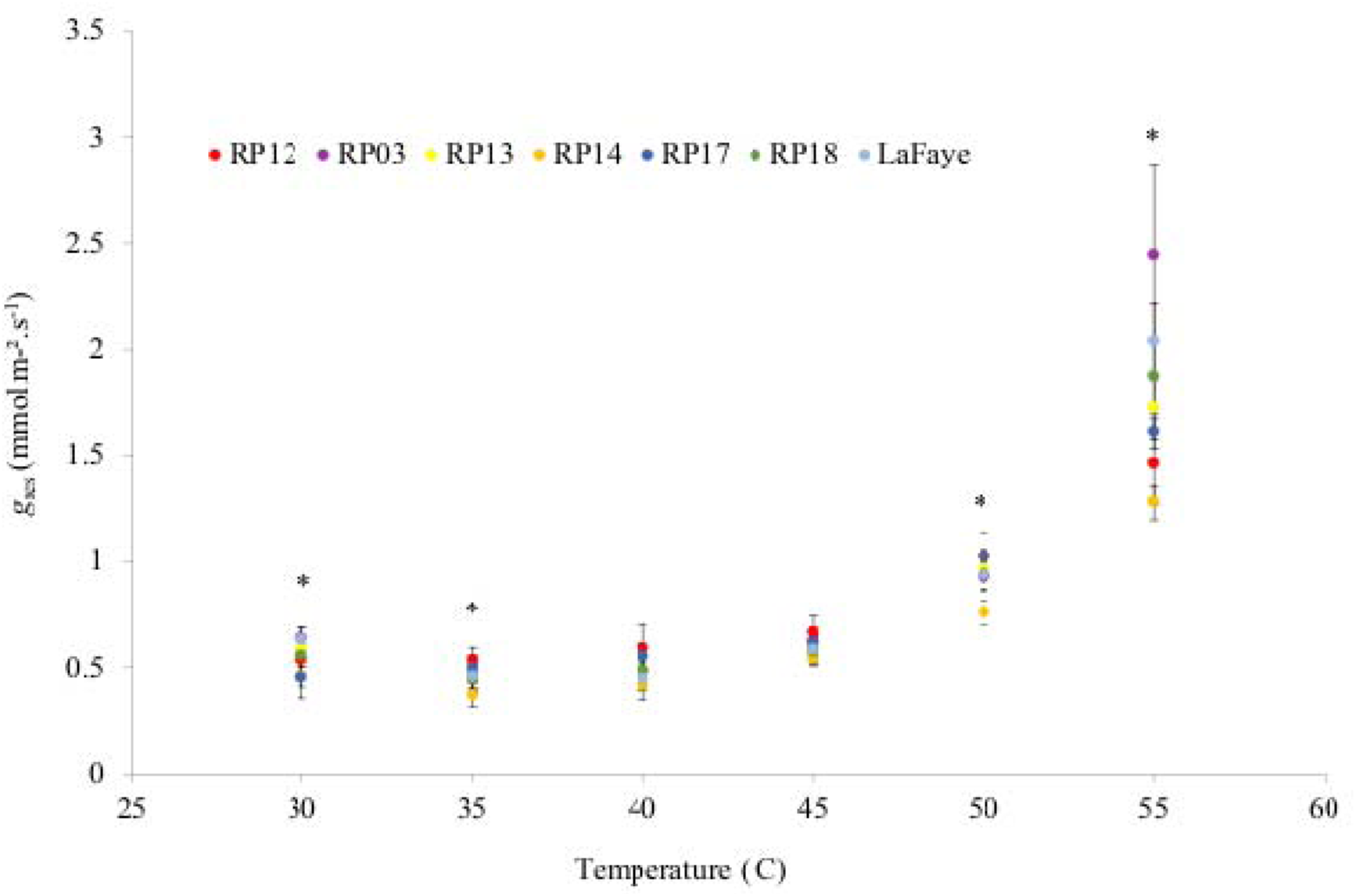
Variability of *g*_res_ measured at different temperatures across seven French fir provenances grown in the same common garden. *g*_res_ was measured at different temperature for the different provenances. These data allow calculating values for *T*_p_, *Q*_10a_ and *Q*_10b_ in the table 2. Data are the mean values (± S.E.). Asterisk indicate significant difference found between at least two provenances at each temperature.

**Table 2.** Hydraulic safety traits for *A. alba* provenances grown in a common garden. Different traits were measured on the same sunlight exposed branches on *A. alba* provenances. Data are the mean values (± S.E.), and *g*_res_ values were determined at 30 °C. Different letters indicate significant differences between provenances for *g*_res_, while for other traits no significant difference was found between provenances.

| Provenance | $\Psi_{TLP}$ (MPa) | $g_{\text{res}}$ ($\text{mmol.m}^{-2}.\text{s}^{-1}$) | $T_p$ ($^{\circ}\text{C}$) | $Q_{10a}$ | $Q_{10b}$ | $P_{50}$ (MPa) |
| --- | --- | --- | --- | --- | --- | --- |
| RP03 | N.D. | 0.635 (± 0.051)<sup>a</sup> | 45.3 (± 0.22) | 1.34 (± 0.10) | 4.10 (± 0.81) | -3.48 (± 0.05) |
| RP12 | -2.35 (± 0.06) | 0.535 (± 0.061)<sup>abc</sup> | 45.2 (± 0.87) | 1.26 (± 0.09) | 2.20 (± 0.11) | -3.58 (± 0.05) |
| RP13 | -2.07 (± 0.06) | 0.583 (± 0.053)<sup>abc</sup> | 44.0 (± 0.64) | 1.29 (± 0.08) | 3.43 (± 0.73) | -3.50 (± 0.05) |
| RP14 | -2.20 (± 0.10) | 0.458 (± 0.047)<sup>b</sup> | 43.8 (± 2.18) | 1.16 (± 0.03) | 2.83 (± 0.23) | -3.39 (± 0.07) |
| RP17 | -2.15 (± 0.06) | 0.453 (± 0.097)<sup>bc</sup> | 45.4 (± 0.28) | 1.45 (± 0.15) | 2.74 (± 0.34) | -3.52 (± 0.07) |
| RP18 | N.D. | 0.556 (± 0.054)<sup>bc</sup> | 43.8 (± 0.78) | 1.17 (± 0.08) | 3.48 (± 0.59) | -3.58 (± 0.07) |
| La Fage | N.D. | 0.633 (± 0.053)<sup>ac</sup> | 44.4 (± 1.04) | 1.17 (± 0.12) | 4.08 (± 0.49) | -3.47 (± 0.05) |

### Environmental plasticity of *g*_res_ and hydraulic safety traits

Residual conductance differed significantly among forest sites along the environmental gradient (Fig. 3). Trees growing at the most favorable sites (Fournols and SJA) exhibited higher *g*_res_ values than those from the less favorable sites (Ceyrat and Condat; Fig. 3). At 30 °C, mean *g*_res_ ranged from 0.891 to 1.425 mmol m=:J² s=:J¹ (Table 3). In contrast, *T*_p_ and *Q*_10a_ and *Q*_10b_ varied little among sites. Vulnerability to xylem embolism (*P*_50_) exhibited only minor variation among sites, with overlapping values across sites.

**Fig. 3.**
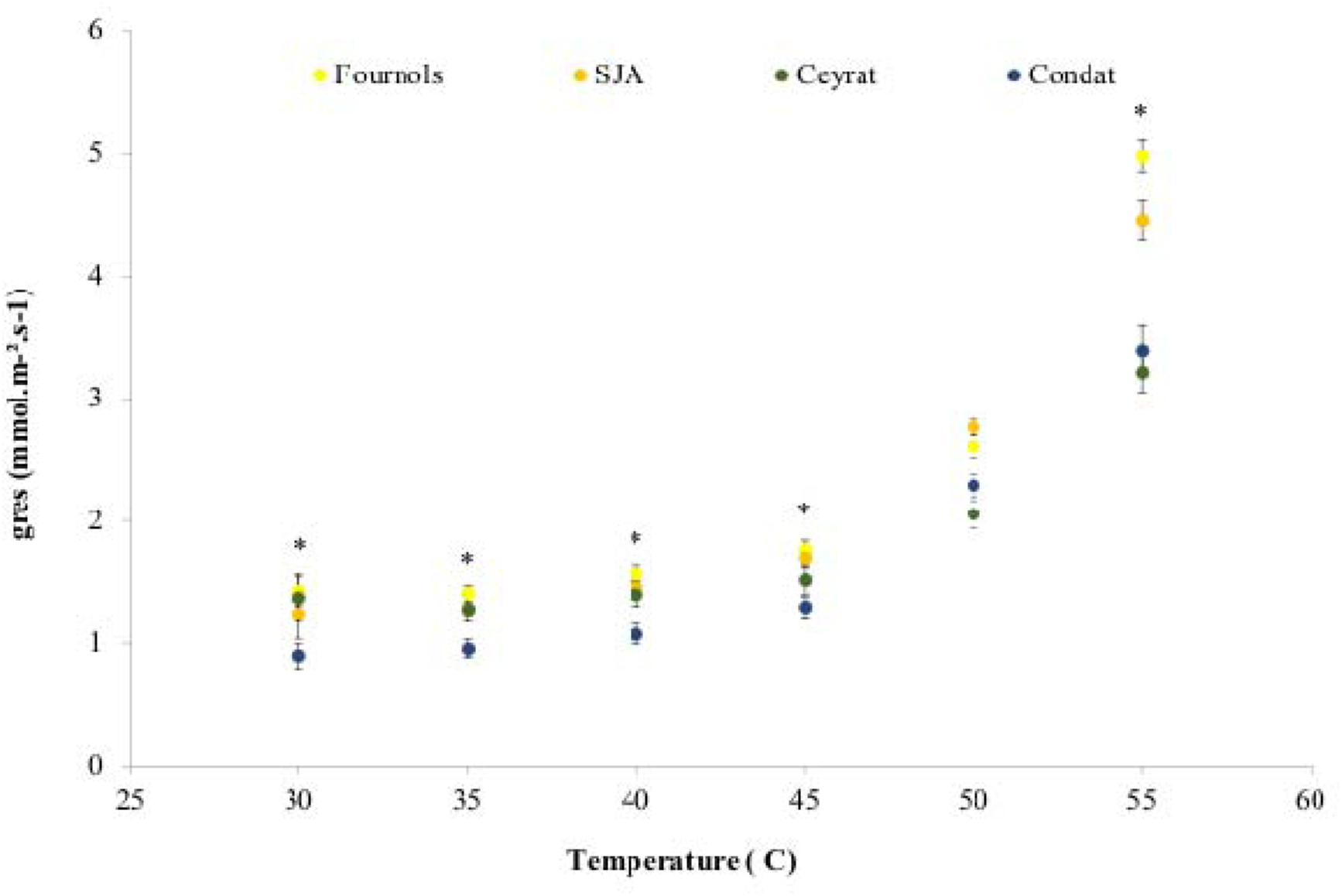
Variability of *g*_res_ measured at different temperatures across Fir trees grown in different forest sites. *g*_res_ was measured at different temperature for the different forest sites. These data allow calculating values for *T*_p_, *Q*_10a_ and *Q*_10b_ in the table 3. Data are the mean values (± S.E.). Asterisk indicate significant difference found between at least two forest sites at each temperature.

**Table 3.**
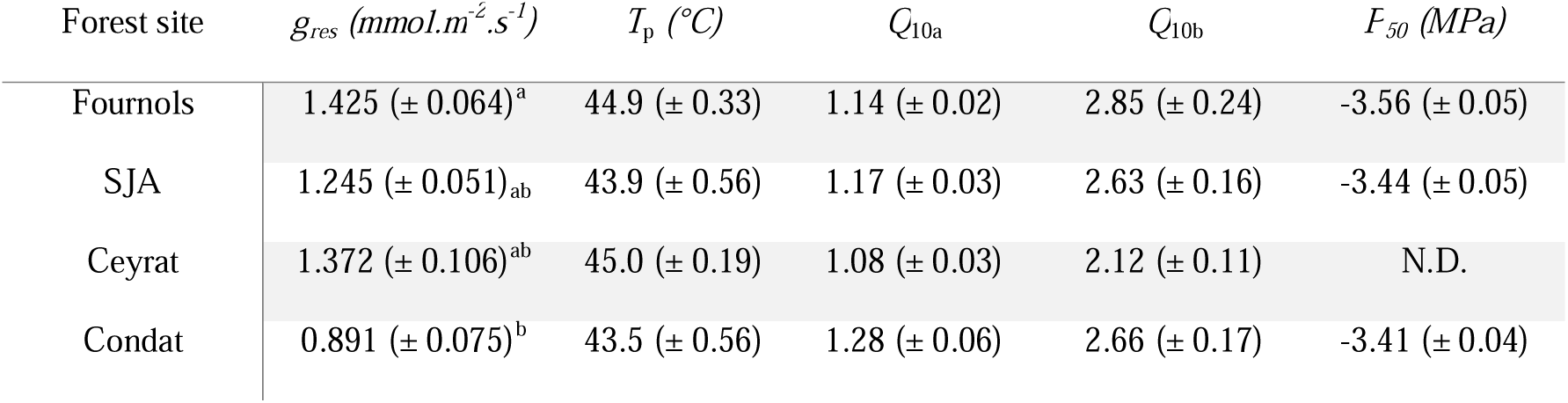
Hydraulic safety traits for Fir grown in different forest sites. Different traits were measured on the same sunlight exposed branches on *A. alba* trees. Data are the mean values (± S.E.), and *g*_res_ values were determined at 30 °C. Different letters indicate significant differences between provenances for *g*_res_, while for other traits no significant difference was found between provenances.

Similarly, no significant differences were detected between sun-exposed upper branches and shade-exposed lower branches for *g*_res_, *T*_p_, *Q*_10a_, *Q*_10b_ or *P*_50_ (Table 4). Mean values for all hydraulic traits were comparable between canopy positions.

**Table 4:** Comparison of hydraulic safety traits between sun-exposed and shade-exposed branches. *P*_50_, *g*_res_, *T*_P_, *Q*_10a_ and *Q*_10b_ were measured on sun-exposed upper branches and shade-exposed lower branches of *A. alba* individuals. Data are mean values (± S.E.). Significant difference were found between *g*_res_ values with *p*-value < 0.05, while no significant differences were found for the other traits.

| Condition | $g_{res} (mmol.m^{-2}.s^{-1})$ | $T_p (^{\circ}C)$ | $Q_{10a}$ | $Q_{10b}$ | $P_{50} (MPa)$ |
| --- | --- | --- | --- | --- | --- |
| Sun-exposed | 0.554 ($\pm 0.063$) | 44.0 ($\pm 0.81$) | 1.31 ($\pm 0.10$) | 3.23 ($\pm 0.87$) | -3.50 ($\pm 0.05$) |
| Shade-exposed | 0.652 ($\pm 0.058$) | 44.1 ($\pm 0.38$) | 1.24 ($\pm 0.16$) | 2.31 ($\pm 0.21$) | -3.49 ($\pm 0.05$) |

### Seasonal variation in needle biochemical parameters across season

Cuticular wax mass exhibited a seasonal variation (Fig. 4a). It was lowest early in the growing season and increased progressively thereafter, reaching higher values during autumn and winter, which were maintained into the following spring on the same cohort of needles. In contrast, Klason lignin content showed more limited seasonal variation, with no clear directional trend (Fig. 4b).

**Fig. 4.**
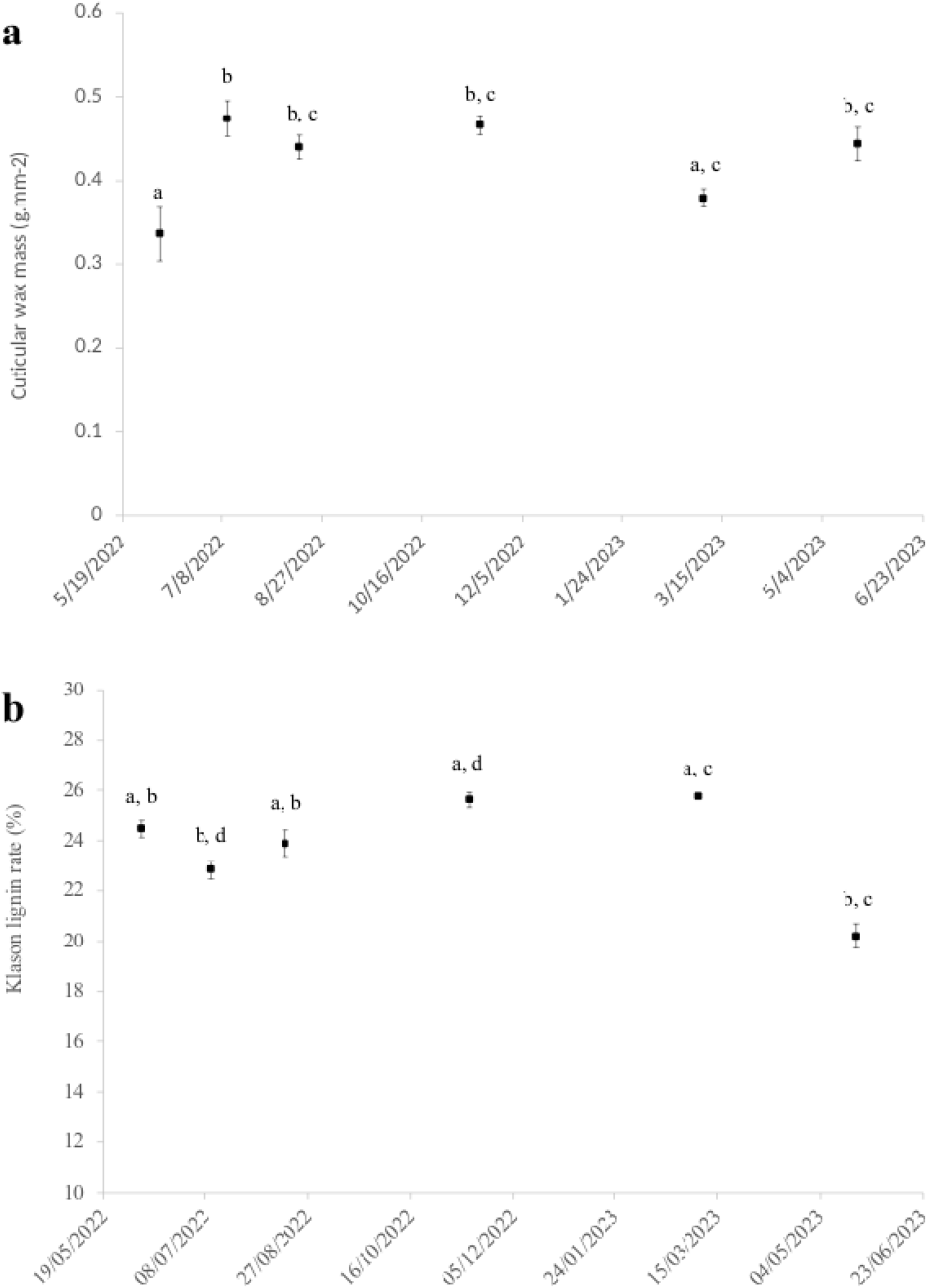
Evolution of needle biochemical parameters across season. Measurements carried out on the same samples as figure 1. Cuticular wax mass (a) and Klason lignin rate (b) were determined as described in the methods section. The individual trees monitored are the same, as is the cohort of needles. The 06/07/2022 measurements are taken on the most recent, fully developed needles, while those taken on 05/ 22/2023 are 1 year old. Data are the mean values (± S.E.). Different letters indicate significant differences between sampling dates at p-value < 0.05.

## Discussion

### 1. Seasonal dynamics of *g*_res_: methodological and functional implications

This study reveals a seasonal pattern in *g*_res_ in *Abies alba*, with high values in newly formed needles in the growing season, followed by a rapid decline and a prolonged stable phase from mid-summer to the following spring. This trajectory highlights the strong dependence of *g*_res_ on needle developmental stage, with important methodological and functional implications. Although seasonal variation in minimum conductance has been reported in some species (Kerstiens 1996; Bueno et al. 2019), it remains less documented than changes in stomatal behavior or xylem vulnerability (Gao et al. 2005; Herbette et al. 2010; Foster et al. 2013). In evergreen conifers, where needle cohorts persist for many years and physiological traits differ between age classes (Oluborode et al. 2025; Ye et al. 2026), failing to control for needle cohort can introduce substantial bias when comparing *g*_res_ across populations, genotypes, or environments - an issue particularly critical for phenotyping studies.

Functionally, the high *g*_res_ of young needles likely reflects incomplete development of diffusion barriers, especially the cuticle, which is initially thinner, less structured, and poorer in epicuticular waxes (Riederer and Schreiber 2001; Kerstiens 2006). Similar developmental patterns have been reported in broadleaved species (Riederer and Schreiber 2001; Burghardt and Riederer 2003), and our data extend these findings to evergreen conifers, showing that ontogenetic effects persist long after full needle expansion. The stabilization of *g*_res_ from late summer through winter indicates that, once the structural barriers are fully developed, residual conductance becomes a conservative trait. This stability parallels that of xylem vulnerability to embolism (*P*_50_, Herbette et al. 2010), suggesting that key hydraulic safety traits stabilized after a critical developmental window. Methodologically, these results argue for standardizing the timing of *g*_res_ measurements. Because *g*_res_ influences time to hydraulic failure in mechanistic models (Cochard 2021), measurements should target ecologically relevant periods, i.e. during summer droughts and heatwaves.

Consistent with the ontogenetic effect of *g*_res_ described above, its seasonal decline closely paralleled the progressive increase in cuticular wax content from early summer to winter, suggesting that wax accumulation is a major driver of this pattern. Cuticular waxes form a hydrophobic barrier that limits water diffusion, and higher wax loads are generally associated with lower cuticular permeability and improved resistance to desiccation (Kerstiens 1996; Riederer and Schreiber 2001; Schuster et al. 2017). Beyond quantity, changes in wax composition and organization can strongly affect permeability (Burghardt and Riederer 2003; Bueno et al. 2019). By contrast, Klason lignin content showed only weak seasonal variation and no clear relationship with gres, indicating that lignification is not a primary determinant of residual conductance in *A. alba*. Although lignification has been linked to drought resistance at the whole-leaf or whole-plant level through increased tissue rigidity and structural integrity (Tu et al. 2020; Choi et al. 2023), our results suggest that diffusion resistance in this species is mainly governed by cuticular properties rather than by internal structural barriers. This is consistent with previous evidence that the cuticle represents the main resistance to water vapor diffusion once stomata are closed (Schuster et al. 2017). Subtle anatomical changes may still contribute indirectly, but these effects are not captured by bulk lignin measurements.

### 2. Limited genetic differentiation among provenances

A main result of this study is the weak difference among provenances for *g*_res_ and its thermal sensitivity (related to *T*_p_, *Q*_10a_, *Q*_10b_), as well as for vulnerability to xylem embolism (*P*_50_).

This is consistent with previous works showing that genetic variation in key hydraulic safety traits is often limited in long-lived forest trees (Wortemann et al. 2011; Lamy et al. 2014; Jinagool et al. 2015, NaN/NaN/NaN; Hajek et al. 2016). In conifers, P50 in particular typically shows low inter-population variability even across wide climatic gradients (Martínez-Vilalta et al. 2009; Lamy et al. 2014), a pattern commonly attributed to strong stabilizing selection or genetic canalization of survival-critical traits (Waddington 1942; Queitsch et al. 2002). The hydraulic safety is a likely target of such canalization (Lamy et al. 2014), as increased vulnerability would incur high mortality costs during drought. Our results suggest that *g*_res_ and its thermal sensitivity may be constrained in a similar way, which is notable given their recently recognized role in controlling time to hydraulic failure (Martin-StPaul et al. 2017; Blackman et al. 2023). If *g*_res_ is under strong selective pressure, its limited genetic variability would be consistent with its functional importance. Accordingly, across multiple *Abies* species, *g*_res_ is the trait that best predicts the time to hydraulic failure, underscoring its central role in drought resistance (Copie et al. 2025). From an applied perspective, the weak provenance differentiation for *g*_res_ and *P*_50_ suggests that selecting material from warmer or drier regions would not necessarily improve drought resistance through these traits alone, as previously reported for xylem vulnerability (Wortemann et al. 2011; Lamy et al. 2014). Actually, forest trees often harbor more genetic variation within than among populations (Hamrick et al. 1992; Petit and Hampe 2006), but our provenance-based sampling may not capture the full extent of intraspecific variability. Adaptive potential may rely more on phenotypic plasticity or on other traits not assessed here and that would represent a tricky task, such as leaf area, rooting depth, or stomatal regulation. In contrast, substantial interspecific differences among fir species have been reported in *g*_res_ and in simulated hydraulic vulnerability, suggesting that adaptation to dry conditions may be more pronounced among species than within a given species (Copie et al. 2025).

### 3. Environmental plasticity of *g*_res_ and associated traits

In contrast to weak differences between fir provenance, *g*_res_ exhibited differences between the four forest sites. Trees at more favorable sites (higher altitude, precipitation, or radial growth) exhibited higher *g*_res_ than those at constrained sites, indicating that *g*_res_ acclimates to drier conditions. While plasticity has been widely documented for other hydraulic safety traits such as stomatal conductance or xylem vulnerability (Awad et al. 2010; Herbette et al. 2010; Lamy et al. 2014; Lemaire et al. 2021; Andriantelomanana et al. 2024), much less is known about intraspecific variation in *g*_res_. Our results show that *g*_res_ can vary substantially within a species in response to environmental conditions, highlighting its potential role in hydraulic adjustment. Lower *g*_res_ at constrained sites likely reflects the development of more efficient diffusion barriers, potentially through changes in wax deposition, cuticle thickness, or composition, consistent with the general expectation that trees in drier or hotter environments reduce non-stomatal water losses (Kerstiens 1996; Schreiber and Schönherr 2009). Surprisingly, the *g*_res_-related thermal parameters, *T*_p_, *Q*_10a_ and *Q*_10b_, remained relatively stable across sites, suggesting that baseline cuticular permeability can be adjusted while thermodynamic properties of waxes remain constrained. To our knowledge, this study is the first to investigate the variability of these thermal parameters in relation to environmental conditions. Further studies across a broader range of species are now required to determine whether the apparent stability of these parameters reflects a general constraint of cuticular functioning or a species-specific feature of silver fir. Vulnerability to xylem embolism (*P*_50_) also showed little variation across sites despite large differences in growth, consistent with previous observations in conifers (Martínez-Vilalta et al. 2009; Barigah et al. 2023). Indeed, a meta-analysis confirmed that the plasticity for hydraulic traits including *P*_50_ is larger in angiosperms than gymnosperms (Anderegg 2015). Because *P*_50_ remained relatively conserved across environmental conditions, the plasticity observed in *g*_res_ likely represents the main hydraulic lever for acclimation to drier environments.

### 4. Implications for drought-induced mortality and heatwave vulnerability

Residual conductance has recently emerged as a key determinant of drought-induced mortality, particularly under conditions of high atmospheric demand when stomata are closed (Blackman et al. 2016; Martin-StPaul et al. 2017; Cochard 2021). Mechanistic models such as SurEau show that even small changes in *g*=:J=:J=:J can strongly affect the time to hydraulic failure by controlling residual water losses during prolonged drought. Simulations with SurEau further indicate that, among some measured traits, *g*_res_ is the primary determinant of the time to hydraulic failure in *Abies* species, highlighting that interspecific differences in drought vulnerability are largely driven by this trait. While *P*_50_ (or *P*_88_) defines the critical threshold for hydraulic failure, *g*_res_ largely governs how fast this threshold is reached (Martin-StPaul et al. 2017; Cochard 2021). Plastic adjustments in *g*_res_ may therefore play a central role in delaying hydraulic failure by reducing dehydration rates and prolonging survival under atmospheric drought.

The thermal tolerance of the cuticle in silver fir is relatively high, with *T*_p_ values ranging from 43.5 to 45.4=:J°C, providing baseline protection against temperature-driven increases in residual conductance. The thermal sensitivity parameters *Q*_10a_ (1.08–1.45) and *Q*_10b_ (2.12–4.10) remain poorly documented, but their moderate values for silver fir indicate that, below Tp, *g*_res_ is unlikely to be strongly affected by temperature. Once *T*_p_ is exceeded, however, *g*_res_ could rise sharply, particularly under high vapor pressure deficits, making residual water losses a dominant pathway for dehydration. Under future climate scenarios combining more frequent heatwaves and intense atmospheric drought, this nonlinear response may critically shape survival outcomes.

By revealing *g*_res_ as a plastic and developmentally dynamic trait, our results emphasize its potential role as a key modulator of drought vulnerability under climate change. Incorporating such traits into trait-based and mechanistic frameworks will be essential for improving predictions of forest responses to increasing climatic extremes.

## Supporting information

raw data

## Author Contributions

SH and HC conceived and supervised the study. SH led the experimental design, performed field sampling, conducted P50 measurements, and wrote the manuscript. LM contributed to experimental design, field sampling, growth measurements, P50 measurements, and manuscript preparation. JC performed residual conductance (*g*_res_) measurements and associated thermal analyses, participated in field sampling, and contributed to manuscript preparation. SA contributed to the experimental design of the seasonal monitoring, carried out biochemical analyses, and contributed to manuscript preparation. AG contributed to experimental design, performed statistical analyses, assisted with data interpretation, and contributed to manuscript preparation. CB performed *g*_res_ measurements and pressure–volume curve analyses and participated in field sampling. All authors read and approved the final manuscript.

## Acknowledgements

The authors are grateful to the French National Forestry Office (ONF) for their help in finding experimental sites.

## Data availability

All data generated and analysed during this study are included in this published article [see supplementary information files].

